# The GCTx format and cmap{Py, R, M} packages: resources for the optimized storage and integrated traversal of dense matrices of data and annotations

**DOI:** 10.1101/227041

**Authors:** Oana M. Enache, David L. Lahr, Ted E. Natoli, Lev Litichevskiy, David Wadden, Corey Flynn, Joshua Gould, Jacob K. Asiedu, Rajiv Narayan, Aravind Subramanian

## Abstract

**Motivation:** Computational analysis of datasets generated by treating cells with pharmacological and genetic perturbagens has proven useful for the discovery of functional relationships. Facilitated by technological improvements, perturbational datasets have grown in recent years to include millions of experiments. While initial studies, such as our work on Connectivity Map, used gene expression readouts, recent studies from the NIH LINCS consortium have expanded to a more diverse set of molecular readouts, including proteomic and cell morphological signatures. Sharing these diverse data creates many opportunities for research and discovery, but the unprecedented size of data generated and the complex metadata associated with experiments have also created fundamental technical challenges regarding data storage and cross-assay integration.

**Results:** We present the GCTx file format and a suite of open-source packages for the efficient storage, serialization, and analysis of dense two-dimensional matrices. The utility of this format is not just theoretical; we have extensively used the format in the Connectivity Map to assemble and share massive data sets comprising 1.7 million experiments. We anticipate that the generalizability of the GCTx format, paired with code libraries that we provide, will stimulate wider adoption and lower barriers for integrated cross-assay analysis and algorithm development.

**Availability:** Software packages (available in Matlab, Python, and R) are freely available at https://github.com/cmap

**Supplementary information:** Supplementary information is available at clue.io/code.

## 1. Introduction

Computational analysis of datasets generated by treating a diversity of cells with pharmacological and genetic perturbagens has proven useful for the discovery of functional relationships (Weinstein *et al.*, 1997); (Hughes *et al.*, 2000; Lamb *et al.*, 2006). To enable such discovery, the NIH Common Fund’s Library of Network-Based Cellular Signatures (LINCS) has brought together a multitude of high-dimensional assays--ranging from gene expression to cellular morphology--to characterize the effects of perturbagens on human cells (Keenan, 2017).

Facilitated by technological improvements, perturbational datasets have grown in recent years from thousands to millions of samples. These large-scale, holistic representations of perturbation provide an incredible opportunity for systems-based research on health and disease. However, they simultaneously introduce data management challenges; one such issue is how large quantities of diverse assay outputs are stored and accessed so that researchers seeking to develop new algorithms can efficiently integrate these data into their computational environment. While conceptually similar concerns are also seen in handling DNA sequencing data, perturbation-based functional compendia present their own set of challenges. Raw forms of data are diverse; for instance, high-throughput mRNA measurements use flow cytometry to detect transcript levels (Subramanian *et al.*, 2017), proteomic readouts produce mass spectrometry traces to assess protein phosphorylation status (Abelin *et al.*, 2016), and quantitative data are extracted from microscopy images in morphological profiling (Bray *et al.*, 2016). With increasing awareness of the importance of metadata, consortia such as LINCS have established standards for data deposition (Vempati *et al.*, 2014), yet there is a gap between adoption of a standard for data deposition and facile access for computation during exploratory data analysis.

The first hurdle of data analysis in this context is that once run through pipelines, resulting data need to be organized in a way that is straightforward to incorporate into developers’ tools and pipelines for downstream analyses. Specifically, while each experiment results in new data, the common unit of analysis is a cohort of related experiments, which is conveniently represented as a matrix wherein each column represents an experiment (treatment) and each row a feature measured by the assay utilized. Text-based formats have long been used in gene expression analysis (Eisen *et al.*, 1998), but the dramatically increasing size of matrices has made storage as plain text impractical (See section 2.2). A second hurdle is that, while thousands of perturbagens may be profiled, many forms of analysis start with a particular drug, pathway or gene of interest. Hence, it is vital to be able to identify and stream relevant portions of the data matrix into a developer’s tools without the memory burden of loading all experiments or all features. Third, the power of a compendium approach is that analysis iterates through all relevant experiments (columns) in a matrix so as to discover unexpected correlations observed in data. Therefore, it should be simple to efficiently traverse the entirety of the matrix and compute a metric (e.g., similarity to an external query signature) for every experiment while also keeping track of experimental conditions and quality control metrics. Fourth, having access to rich sets of metadata annotations from the literature on drug targets, mechanisms of action (Corsello *et al.*, 2017) and pathways (Liberzon *et al.*, 2011) is valuable for interpretation of analysis results; furthermore, it is convenient for analysis and helpful for maintaining reproducible workflows when such annotations are packaged together with quantitative data. Finally, as several existing machine learning and bioinformatics toolkits can be readily applied to analyze perturbational data, new data representations should ideally be straightforward to retrofit.

As solutions to these analysis challenges, we present the GCTx file format along with open-source software packages that we have developed. GCTx relies on robust HDF5 technology ({The HDF Group}, 1997-2014) to make large, dense matrices of data and metadata annotations easy to store and explore. Importantly, the format’s utility is not just theoretical: to date, we have compiled 1,677,442 samples and made them freely available in the GCTx format.

## 2. Features

### 2.1 The GCTx format and code libraries

#### 2.1.1 Choice of a data representation technology

HDF5 supports a platform-independent file format capable of unlimited size, a customizable data model, and highly performant read/write capabilities {The HDF Group}, 1997-2014). The format scales with data size by using hyperslabs to selectively parse portions of the matrix, thereby making it possible to load only a subset of interest from a data file that may or may not exceed physical memory limits. HDF5 has a vibrant developer community that supports multiple programming languages and operating systems.

#### 2.1.2 The GCTx Format

Although HDF5-based formats have previously been used in genomics (Millard *et al*., 2011; Sommer *et al*., 2013), they have been tailored towards different use cases than that of GCTx; some are optimized to capture detailed schema on experimental design and modeling complex relationships between entities and data, while others have used HDF5 to bundle heterogeneous data types into a single file. While these are useful attributes, our primary need was a format optimized to handle massive and dense two-dimensional data matrices. To achieve this, we developed a schema built on HDF5, that we term GCTx, which adopts a lightweight, shallow hierarchy for representing matrices and associated annotations while retaining several of the key benefits of the HDF5 infrastructure. GCTx is a more optimized version of a prior text format called GCT (Gene Cluster Text), which stores data alongside metadata in a single tab-delimited text file (Figure 1A).

**Figure 1.**
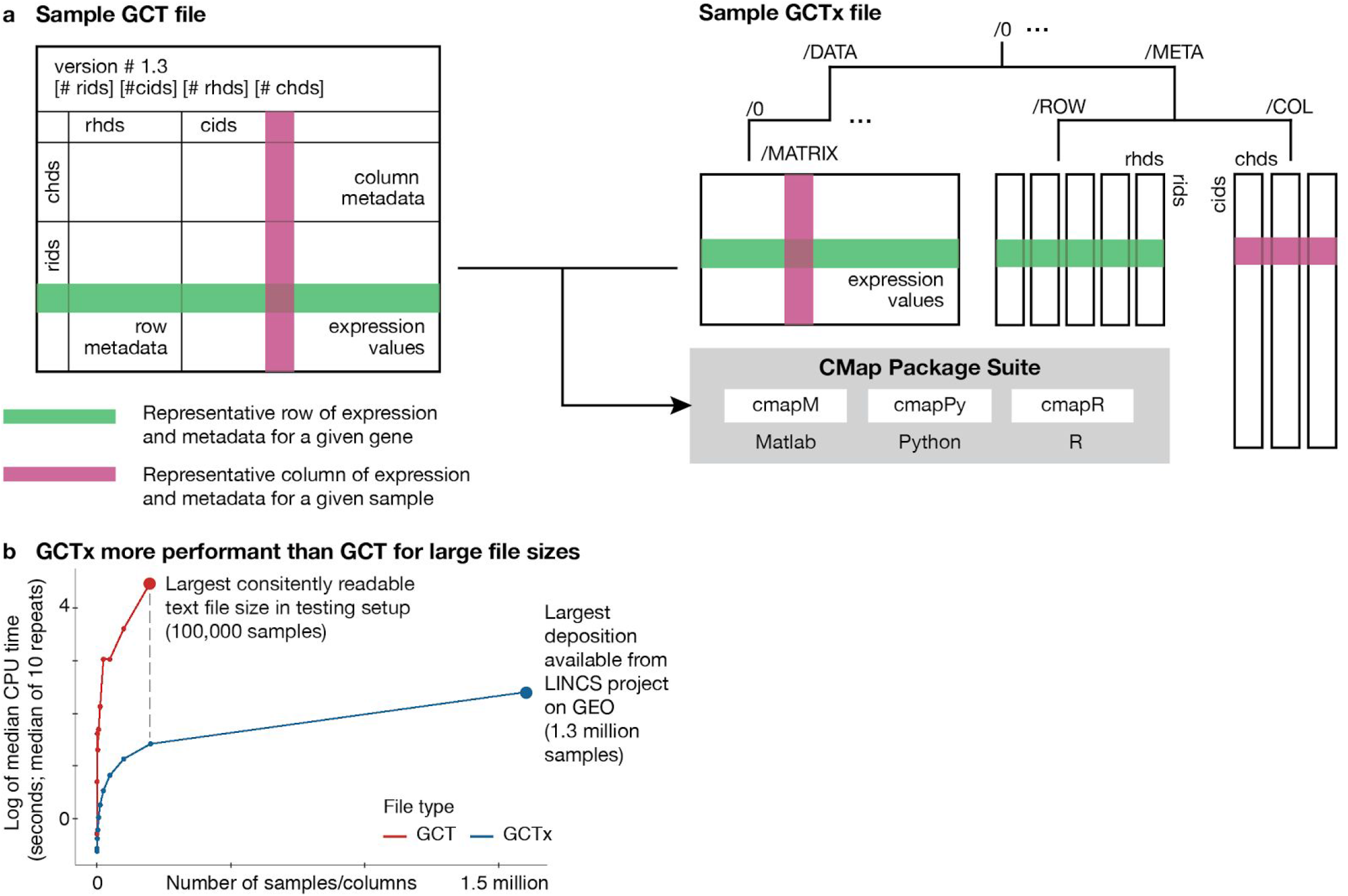
**(a) Schematic of a GCT and GCTx file**. The GCTx format can contain the same content as GCT, but in the hierarchical structure of HDF5 instead of as tab-dθl¡m¡ted text. Note that rids and rhds for rows (or cids and chds for columns) respectively correspond to the unique identifiers of samples and headers for metadata fields. The CMap suite of packages accept either GCT or GCTx files and enable users to easily access and modify these files’ contents, **(b) Parse times are considerably faster for GCTx files compared with GCT, particularly as file size increases**. Parse times for text-based formats grow exponentially as the number of samples included increases, while the time to parse files of equivalent content in GCTx format increases more linearly; for this reason, we were unable to consistently parse files of over 100,000 samples on the EC2 instance provisioned fortesting. This along with GCTx’s compact file sizes enables users to also parse more samples into memory than with text-based formats. For example, medium CPU time to parse the largest LINCS file published to GEO (accession GSE 92742) with methods from cmapPy took 252.2 seconds on testing setup.

The GCTx format offers a number of benefits: matrix lookups might need to serially iterate through all columns, to access logically contiguous collections of columns, or to access random subsets of columns/rows. GCTx is performant for all these forms of access, freeing a developer from having to repeatedly customize their analytical code. By optimizing the schema for matrices, the task of analysis is simplified; users do not have to think about how to optimally use or represent the data in HDF5. This is important because HDF5 only provides a generic data model; consequently, standardizing the representation of data and metadata encourages reuse. GCTx preserves the high performance and I/O performance benefits of HDF5, and its shallow hierarchy of component nodes decreases random access time when compared to deeply nested alternatives.

An important design consideration for the format was to enable support for storing multiple datasets based on common data usage patterns and computational workflows. To that end, the GCTx format is extensible to support two specific use cases. First, when a dataset is subject to transformations as part of a data processing pipeline (e.g. raw, normalized or re-scaled), the dimensionality of the underlying matrices frequently remains the same, as does the associated metadata. Storing these data at each level of processing in a single file is enabled by simply adding additional matrices of the same dimension to the /MATRIX node after incrementing the numerically indexed group name (“/0”, “/1”, and so on). This layout has the additional benefit of avoiding duplication of the associated metadata. Second, datasets of variable dimensions and metadata that are encountered in computational workflows can be stored in the same file as separate matrix and metadata nodal entities after incrementing the index of the root group.

#### 2.1.3 Open source software packages integrated with GCTx

To facilitate adoption of the GCTx format by existing bioinformatics and data science tools, we developed three open source software packages in Python, R, and Matlab which simplify input, output, and analysis of GCTx files. Note that the Python package also supplies a command line utility for the conversion of GCT to GCTx files and vice versa. In each of these languages we ensured that GCTx inputs, once processed by the provided code library, are represented as native data structures readily compatible with powerful data analysis and machine learning tools such as the Statistics and Machine Learning Toolbox in Matlab (MathWorks, 2012), pandas and scikit-learn in Python (McKinney, 2012), and the tidyverse package suite in R (Wickham, 2016). Installation instructions, documentation, and tutorials for the Matlab (“cmapM”), Python (“cmapPy”) and R (“cmapR”) packages are provided at clue.io/code; source code is available at github.com/cmap.

#### 2.1.4 Role of GCTx in a cloud-based data commons ecosystem

Relational databases and cloud-based object stores are also capable of storing and efficiently serving massive datasets. Our development of the GCTx schema for internal analysis and public distribution was driven first by user needs and second to maximize accessibility. Databases, while efficient for structured queries, are not optimized for matrix operations. Object stores are highly parallelizable, but impose no structure at all on their contents. In contrast, the GCTx format strikes a good balance between providing highly performant matrix operations while also imposing a model that links to relevant metadata.

Although this may change over time, we have found that even as cloud-based object stores become more commonplace, users still request downloadable file-based representations of data for use on their personal computers or traditional login servers. Prototyping new algorithms is often done in such environments and GCTx and our provided code libraries facilitate this for data matrices of varying sizes. Additionally, it is worth noting that even cloud-based tools often resort to accessing data from local caches to avoid network latency.

### 2.2 GCTx performance benchmarking

To assess the performance of GCTx files compared to their text file equivalents, we performed benchmarking tests that parsed various sizes of text (GCT) and GCTx files of identical content. Benchmarking was performed on an Amazon Web Services EC2 instance with a locally mounted solid state drive (instancetype = i3.16xlarge, memory = 488 GB, vCPU = 64). For each of the software packages available (cmapM, cmapPy, cmapR), the same set of representative files of expression data was parsed 10 times (cache cleared between sequential reads). The median times for all parsing operations performed in Python are plotted in Figure 1B; similar trends were observed for the Matlab and R implementations. The text-based GCT files become considerably slower to read from and write to as file sizes increase beyond tens of thousands of columns/samples, and were not performant on the testing setup used as file sizes increased beyond a hundred thousand samples/columns. However, GCTx can parse and write equivalent and much larger content considerably faster, and continues to be performant for much larger file sizes.

### 2.3 Publicly available data sets using GCTx

GCTx’s capability to portably represent diverse data types has lead to its adoption by many projects; currently, publicly available data in GCTx format include L1000 gene expression datasets from the Connectivity Map (GSE70138/GSE92742), phosphorylation and chromatin datasets from LINCS (GSE101406; Litichevskiy *et al.*, 2017), and RNA-Seq datasets from the Cancer Cell Line Encyclopedia (Barretina *et al.*, 2012) and GTEx (Carithers *et al.*, 2015).

## 3. Conclusions

We present GCTx, an HDF5-based file format designed for the space-efficient storage and rapid access of dense data matrices paired with metadata annotations. The format’s ability to store multiple distinct data sets and their annotations enable a single file to contain an entire analysis workflow’s worth of content, which additionally aids reproducibility in analyses and collaboration. To make it easier for users to adopt GCTx, we also present three open-source packages--cmapM, cmapPy, and cmapR--that make GCTx straightforward to incorporate with existing bioinformatic analysis frameworks.

## Acknowledgements

We thank Jodi Hirschman for technical editing of the manuscript and helpful conversations, and Wen Niuw for thorough feedback on software packages and the text. We are grateful to all our users and contributors for using, testing, and providing valuable feedback and additions to both the data and software throughout the development process. We also thank the LINCS consortium for data and the LINCS Data Coordination and Integration Center for user testing and input on metadata stored.

